# Critical scaling of cortical ensembles without cooperative amplification

**DOI:** 10.64898/2026.10.07.757392

**Authors:** Tiago L. Ribeiro, Raiyyan Siddiqui, Bridgette Gifford, Sinisa Pajevic, Dietmar Plenz

**Affiliations:** Section on Critical Brain Dynamics, National Institute of Mental Health, Bethesda, MD, USA

**Author notes:** Correspondence, Dietmar Plenz, Ph.D., Section on Critical Brain Dynamics, National Institute of Mental Health, Porter Neuroscience Research Center, Rm 3A-1000, 35 Convent Drive, Bethesda, MD 20892.

**Keywords:** Perturbation, ensembles, population coding, criticality, scaling, neuronal avalanches, visual cortex, in vivo, 2-photon imaging, holography, all-optical

## Abstract

Cortical neurons are sparsely and weakly connected, motivating the long-standing idea that they must act collectively, as cell assemblies, to exert substantial influence on surrounding networks. Yet a few spikes in a single pyramidal neuron can recruit thousands of cortical neurons according to a power law with exponent ∼0.285, consistent with a two-dimensional system near a critical percolation transition. Here we holographically stimulated ensembles of 1–16 layer 2/3 pyramidal neurons in visual cortex of awake mice while imaging hundreds of neighbouring neurons. Across two orders of magnitude in evoked spike count, network responses followed the same scaling law observed for single-neuron perturbations, with no detectable threshold at which co-activation produced additional amplification. Recruitment was sublinear, suppression increased with ensemble size, and ongoing scale-free avalanche dynamics remained stationary. Nevertheless, ensemble identity became increasingly decodable from surrounding activity. Thus, coordinated firing increases representational specificity without escaping the critical dynamics governing propagation: cortical ensembles convey identity without gaining disproportionate causal impact.

## Introduction

Cortical computation is commonly thought to depend on groups of neurons firing together. This idea has deep roots in Hebbian cell assemblies and is reinforced by the sparse, weak and failure-prone synaptic connections between neighbouring pyramidal neurons in superficial cortex [1–7]: individual neurons should exert little influence, whereas coincident activity from multiple neurons can converge to reliably drive their targets. Variants of this principle underlie models of synfire chains [8,9], travelling waves [10,11], phase-locked communication [12,13] and, more generally, neuronal ensembles whose coordinated activity is amplified as it propagates through cortical networks [6,14]. Two-photon holographic optogenetics has provided striking support for the functional importance of such ensembles. Activating only a few tens of layer 2/3 (L2/3) pyramidal neurons can recall learned representations or behaviour through pattern completion [15,16], and stimulation of as few as 14–20 neurons can influence perception [15,17]. Yet a fundamental assumption underlying these observations remains largely untested: does an ensemble acquire collective influence beyond that expected from simply adding the spikes of its constituent neurons?

Our recent findings suggest a different possibility. Evoking only a few additional spikes in a single L2/3 pyramidal neuron in awake mouse visual cortex produced a widespread transient response involving thousands of surrounding neurons [18]. Approximately 15% of imaged neighbours increased their firing within 132 ms, and the magnitude of recruitment scaled with the number of evoked spikes according to a power law with an exponent of 0.2–0.3. The identity of the stimulated neuron could also be decoded from the resulting population response. These perturbations occurred against ongoing activity organized into scale-invariant, parabolic neuronal avalanches [19–24]. A two-dimensional network poised near criticality reproduced both the distributed propagation of single-neuron perturbations and their scaling exponent, close to the value of 0.285 associated with directed percolation in two dimensions [25–27]. Thus, even sparse local input can propagate widely when the surrounding network operates near a critical point.

This framework leads to a prediction that differs fundamentally from cooperative models of ensemble propagation. Near criticality, the response to a local perturbation scales as a power law of input strength with an exponent below one [25,26,28]. Because this scaling is determined by the dynamical state of the network, increasing the number of spikes should move the response along the same compressive input–output function, regardless of whether those spikes originate from one neuron or are distributed across several co-active neurons. In contrast, models that assign special importance to coincident ensemble activity predict that sufficiently synchronous inputs should recruit shared postsynaptic targets cooperatively, producing supralinear amplification or a transition to a distinct regime of network propagation [8,9,29]. Distinguishing these possibilities requires varying ensemble size while measuring both the imposed spike input and the population response within the same cortical network.

Here we holographically stimulated hierarchically nested ensembles of 1, 2, 4, 8 and 16 L2/3 pyramidal neurons in primary visual cortex of awake mice while monitoring hundreds of neighbouring neurons with two-photon calcium imaging. Across two orders of magnitude in evoked spike count, the population response remained on a single power law with an exponent of approximately 0.285, continuous with that observed for single-neuron perturbations. We detected no ensemble size at which co-activation produced disproportionate amplification; instead, recruitment grew sublinearly and suppression increased with ensemble size, while ongoing avalanche dynamics remained stationary. Yet larger ensembles generated progressively more distinguishable population responses: ensemble identity could be decoded with increasing accuracy and robustness, largely independently of global network activity, with errors concentrated among targets separated by less than approximately 100 μm. Thus, co-active cortical neurons gain representational specificity without acquiring disproportionate causal influence. Near criticality, cortical networks scale ensemble input rather than transforming it into a qualitatively different mode of propagation.

## Results

### Holographic stimulation of hierarchically grouped ensembles of L2/3 pyramidal neurons in awake mice

We tested how cortical responses change as increasing numbers of neurons are activated together. In awake, quietly resting mice expressing jGCaMP8s and ChrimsonR in L2/3 pyramidal neurons, we selected 32 target cells in each V1 field of view (450 × 450 µm; ∼80–160 µm depth). We stimulated these cells holographically for 100 ms (∼5–10 mW per target) while imaging the spiking activity of ∼100–300 surrounding neurons at 22-ms resolution (Fig. 1a,b).

**Fig. 1.**
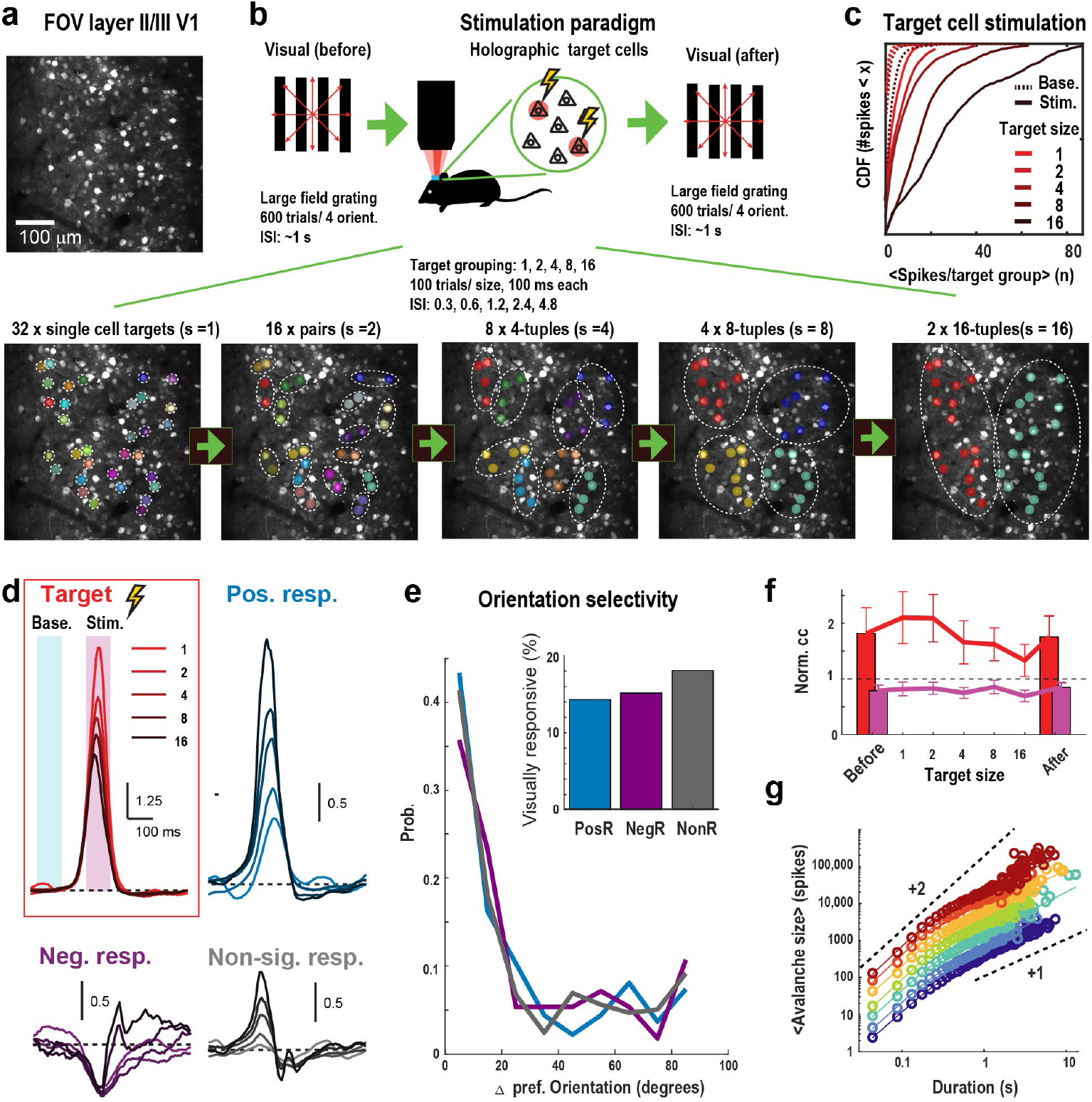
Holographic stimulation of hierarchically grouped pyramidal-neuron ensembles in L2/3 of awake mouse V1. **a,** Mean two-photon image of a 450 × 450 µm field of view in L2/3 V1. **b,** Experimental paradigm. Large-field drifting gratings were presented before and after holographic stimulation to determine orientation preference and record ongoing activity. Thirty-two target cells were stimulated for 100 ms in 100 semi-randomized trials per group, first individually (s = 1) and then as hierarchically merged ensembles of 2, 4, 8 and 16 neurons, with inter-stimulus intervals of 0.3, 0.6, 1.2, 2.4 and 4.8 s, respectively. Each ensemble size was recorded separately in the same field of view. Bottom, example hierarchical grouping of targets; dashed outlines denote ensembles and colours denote groups. **c,** Cumulative distributions of target-group spike counts during stimulation (Stim., solid) and matched baseline windows (Base., dotted) for ensemble sizes 1–16. **d,** Mean baseline-normalized spike-rate time courses for targets, positive responders (PosR), negative responders (NegR) and non-significant responders (NonR). Shading denotes baseline and stimulation windows. Scale bars, indicated normalized spike rate and 100 ms. **e,** Distribution of differences in preferred orientation between target neurons and their corresponding non-targets for PosR (blue), NegR (purple) and NonR (grey). Inset, fraction of visually responsive neurons in each class. **f,** Normalized pairwise correlations across recordings before, during and after ensemble stimulation for target–target pairs (red) and targets versus the remaining population (magenta). Error bars, mean ± s.e.m. across experiments. **g,** Mean neuronal-avalanche size versus duration before (blue), during and after (red) ensemble stimulation. Dashed lines indicate slopes of 2 and 1.

The same 32 target cells were stimulated first individually and then in ensembles of 2, 4, 8 and 16 neurons. Ensembles were constructed hierarchically by grouping spatially neighbouring targets, such that each ensemble was formed by combining two groups from the preceding level (Fig. 1b, bottom). Each ensemble size was tested in a separate ∼17-min recording from the same field of view, with 100 semi-randomized trials per group. Inter-stimulus intervals increased with ensemble size (0.3, 0.6, 1.2, 2.4 and 4.8 s for groups of 1, 2, 4, 8 and 16 neurons, respectively). Before and after the stimulation series, we presented large-field drifting gratings to measure orientation preference and recorded ongoing activity (Fig. 1b, top). The dataset comprised 10 recordings from 9 mice. Beam profiling, motion correction and anatomical analyses were performed as described previously [18].

Holographic stimulation reliably evoked spikes in the targeted neurons. Evoked spike count increased with ensemble size, from a few spikes for single targets to several tens of spikes for 16-cell ensembles (Fig. 1c; Fig. 2a). Our perturbations therefore spanned approximately 1–100 spikes within 100 ms, extending the 1–7-spike range of our previous single-neuron experiments [18] by more than an order of magnitude.

**Fig. 2.**
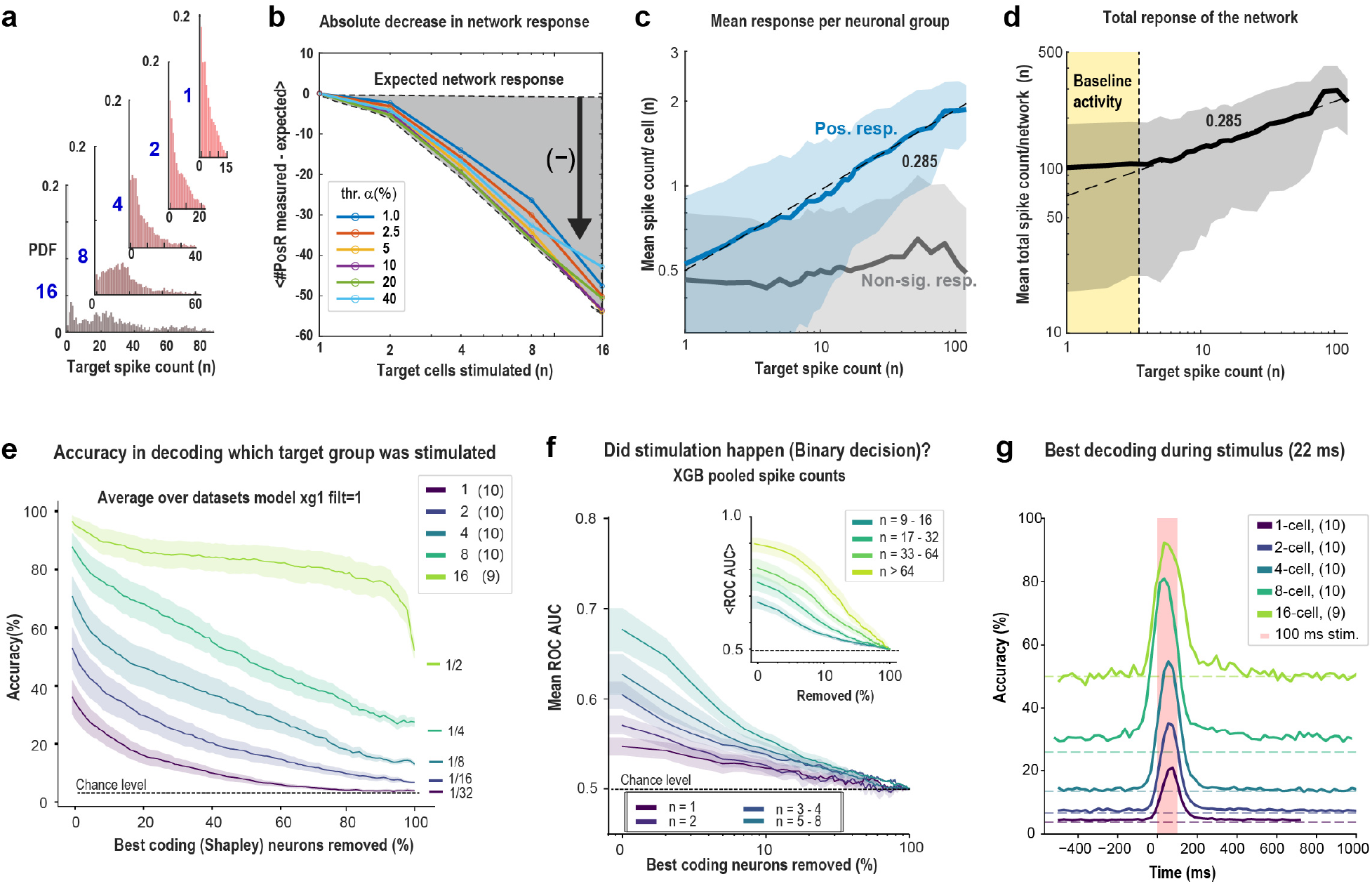
Ensemble responses scale sublinearly, while ensemble identity is distributed across the network. **a,** Distribution of spikes evoked per trial in target ensembles of 1, 2, 4, 8 and 16 neurons; note differing abscissa ranges. **b,** Difference between measured PosR number and that expected from the union of constituent single-target PosR pools, across responder significance thresholds α = 1–40%. Negative values indicate subadditive recruitment. **c,** Mean response-window spike count per neuron versus total evoked target spikes for PosR (blue) and NonR (grey), pooled across ensemble sizes. Dashed line, power law with exponent 0.285; shading, mean ± s.d.. **d,** Mean total non-target spike count versus evoked target spike count. Yellow region marks baseline activity level; dashed line, power-law scaling with exponent 0.285 above the baseline floor. Shading, mean ± s.d.. **e,** XGBoost decoding accuracy for ensemble identity as increasing fractions of the highest-ranked neurons, defined by Shapley values, are removed. Curves show ensemble sizes 1–16 (32, 16, 8, 4 and 2 classes; n = 10 datasets, n = 9 for 16-cell ensembles). Chance levels are indicated at right; shading, mean ± s.e.m.. Trial-filtered data shown; controls are in Supplementary Fig. 2c. **f,** Area under the ROC curve for binary classification of stimulation versus baseline as increasing fractions of the highest-ranked neurons are removed. Trials are grouped by total evoked target spikes (1, 2, 3–4 and 5–8; inset, 9–16, 17–32, 33–64 and >64). Shading, mean ± s.e.m.. **g,** Time-resolved decoding of ensemble identity from single 22-ms frames. Dashed lines, chance; pink shading, 100-ms stimulation period.

As observed for single-neuron perturbations [18], ensemble stimulation produced a widespread but transient response in the surrounding network. Non-targeted neurons were classified as positive responders (PosR), negative responders (NegR) or non-significant responders (NonR) according to whether their firing significantly increased, decreased or remained unchanged during stimulation (α = 2.5%, z-test; Methods). PosR activity followed the time course of the stimulated targets and returned to baseline within ∼100 ms after stimulus offset (Fig. 1d). NonR neurons showed a brief increase followed by suppression below baseline, and this biphasic response became more pronounced with increasing ensemble size (Fig. 1d, bottom right). Thus, larger ensembles increased both recruitment and suppression in the surrounding network. Consistent with this, the fraction of neurons classified as PosR increased with ensemble size across distances from the stimulated targets (Fig. 3a).

**Fig. 3.**
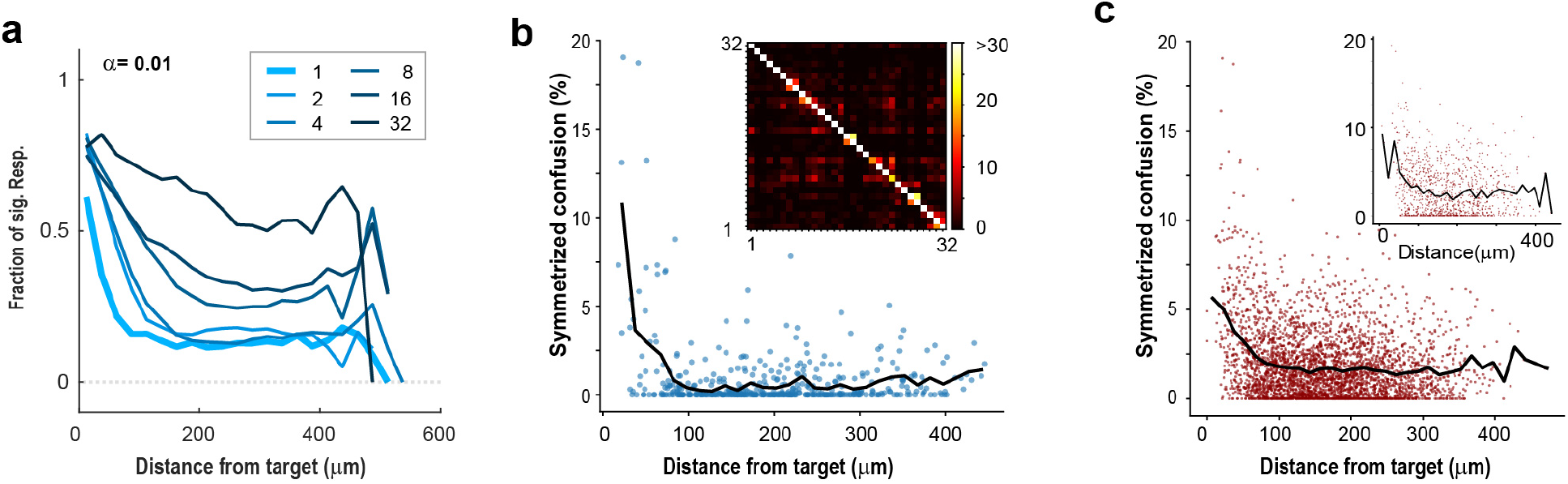
Responder recruitment and decoding confusion are spatially organized around stimulated targets. **a,** Fraction of significant responders (α = 1%) versus distance from ensemble targets for ensemble sizes 1–32. Profiles extend to the field-of-view boundary. Other significance thresholds and responder classes are shown in Supplementary Fig. 5. **b,** Pairwise decoding confusion versus distance between single targets in an example dataset (32 targets). Dots, target pairs; line, mean in ∼13-µm distance bins. Off-diagonal entries of the row-normalized confusion matrix were symmetrized by harmonic mean. Inset, full 32 × 32 confusion matrix; colour scale saturated above 30%. **c,** Pairwise decoding confusion versus target distance pooled across datasets. Inset, corresponding analysis for 2-cell ensembles using centre-of-mass distance. Linear regression analysis: 1-target ensembles (3564 pairs over 10 datasets) – slope −0.00377 %/µm; intercept 3.33%; r −0.187; p(r) 2×10^-9^; ρ −0.158; p (ρ) 2.8×10^-21^; 2-target ensembles (1082 pairs over 10 datasets) – slope −0.00449 %/µm; intercept 4.56%; r −0.151; p(r) 6.3×10^-7^; ρ −0.123; p (ρ) 5×10^-5^ (p-values are two-sided and Spearman ρ is rank-based and robust to the nonlinearity at short distances).

We next asked whether recruitment depended on functional similarity between stimulated and neighbouring neurons [30]. Targets were selected according to opsin expression and location rather than visual tuning. The difference in preferred orientation between non-target and target neurons was similar for PosR, NegR and NonR populations, and the fraction of visually responsive neurons was comparable across the three groups (∼15–18%; Fig. 1e). Thus, recruitment by ensemble stimulation was not detectably organized by orientation preference.

Repeated ensemble stimulation did not produce persistent changes in ongoing network dynamics. Normalized target–population pairwise correlations were similar before and after the stimulation series, and target-pair correlations changed only modestly across ensemble sizes (Fig. 1f).

We further tested whether stimulation altered the scale-invariant dynamics of ongoing activity. As in our previous single-neuron experiments [18], population activity was organized into neuronal avalanches whose mean size scaled with duration with an exponent of approximately 2 for short avalanches, consistent with parabolic avalanche dynamics [19,22,31], before approaching near-linear scaling for the longest events that extended beyond the imaging field [19,22,32] (Fig. 1g). Avalanche size and duration distributions remained parallel before, during and after stimulation of ensembles of 1, 2, 4, 8 and 16 neurons (Supplementary Fig. 1a,b). Similar scaling exponents (χ ≈ 1.8–2.4) were also obtained when recordings were divided according to high- and low-correlation target pairs (Supplementary Fig. 1c,d).

Thus, repeated ensemble stimulation sampled the network across a wide range of imposed input strengths without measurably shifting its ongoing correlation structure or avalanche regime.

### Network responses scale with ensemble input as a single power law despite sublinear responder recruitment

We next asked how network responses scale with the strength of ensemble input. In our previous single-neuron experiments, the spike count of positive responders (PosR) scaled with target-cell spike count as a power law with exponent ∼0.285 [18]. We therefore tested whether increasing the number of co-active neurons would preserve this relationship or produce a new regime of network recruitment.

Responder recruitment was strongly sub-additive. Because each ensemble was constructed by combining previously stimulated smaller groups, we could estimate the number of PosR expected if each constituent target contributed its own independent responder population. Specifically, we calculated the union of the PosR pools recruited by the corresponding single targets. The measured number of PosR increasingly fell below this expectation as ensemble size grew, with a deficit of tens of neurons for 16-cell ensembles, corresponding to approximately half the expected responders (Fig. 2a,b). This effect was robust across significance thresholds ranging from α = 1–40%. Thus, larger ensembles recruited increasingly overlapping rather than additive responder populations, consistent with compressive network recruitment and with the stronger suppression observed at larger ensemble sizes (Fig. 1d) [33,34].

Despite this sublinear recruitment, the activity of recruited neurons followed a remarkably consistent scaling relationship. Pooling across all ensemble sizes, the mean spike count per PosR increased as a single power law with exponent close to 0.285 as the total number of evoked target spikes increased from ∼1 to ∼100 (Fig. 2c; Supplementary Table 1). Mean PosR activity rose approximately four-fold (∼0.5 to ∼2 spikes per response window) across this range. Non-significant responders (NonR), in contrast, showed no systematic dependence on input strength.

The same scaling was evident at the population level. The mean total spike count of all non-target neurons remained near ongoing baseline levels for inputs below ∼3–4 evoked spikes, but above this floor increased with a power-law exponent of ∼0.285 (Fig. 2d). The exponent measured for ensemble stimulation was indistinguishable from that obtained previously for single-neuron perturbations [18] and from the value of 0.285 predicted for a two-dimensional critical network [25] (Supplementary Fig. 2a,b).

Thus, from one to sixteen co-active neurons and across approximately two orders of magnitude in evoked spike count, cortical responses remained on a single compressive scaling curve. Above the ongoing-activity floor, increasing ensemble size revealed no additional inflection, threshold or transition to supralinear amplification.

### Ensemble identity is encoded across the network with increasing robustness

Because ensembles of different sizes propagated along the same scaling curve, we asked whether the surrounding network nevertheless retained information about which neurons had been stimulated. We trained gradient-boosted decision-tree classifiers (XGBoost [35]) on response-window spike counts from non-target neurons, using the stimulated group as the class label and fitting separate classifiers for each dataset and ensemble size (32, 16, 8, 4 and 2 classes for ensembles of 1, 2, 4, 8 and 16 neurons; Methods). Ensemble identity was decoded well above chance at every size (Fig. 2e). Absolute accuracy increased from ∼37% for single targets to ∼53%, ∼70%, ∼88% and ∼97% for ensembles of 2, 4, 8 and 16 neurons, respectively (chance: 3.1%, 6.3%, 12.5%, 25% and 50%). Thus, larger ensembles generated increasingly distinguishable population responses, while even the identity of a single stimulated neuron remained strongly represented in the surrounding network.

This information was distributed and redundant across the population. We ranked neurons by their contribution to the classifier using Shapley values [36,37], progressively removed the highest-ranking neurons and retrained the classifier after each step. Accuracy declined gradually rather than abruptly (Fig. 2e). For 16-cell ensembles, decoding remained above 80% after removing ∼90% of the highest-ranking neurons, and across ensemble sizes a substantial fraction of the population had to be removed before performance approached chance. Ranking neurons by leave-one-out loss, or including all trials without a responsiveness filter, produced similar results (Supplementary Fig. 2c).

At the level of individual trials, however, classifier evidence was concentrated in relatively few neurons. For each target, the largest Shapley values were associated with a small number of highly selective neurons (Supplementary Fig. 3a,b). For well-decoded targets, these neurons contributed strongly and consistently across trials, producing a low-entropy distribution of classifier evidence. For poorly decoded targets, contributions were more diffusely distributed across the population (Supplementary Fig. 3c,d). Thus, ensemble identity is represented sparsely on individual trials but redundantly across the population: a small subset of neurons carries most of the evidence at any moment, yet many others can support decoding when the strongest contributors are removed.

We next asked whether the network also encoded the occurrence of stimulation, independently of ensemble identity. A binary classifier trained to distinguish stimulation trials from baseline windows became progressively more accurate with the total number of evoked target spikes. The area under the ROC curve increased from ∼0.55 for a single evoked spike to ∼0.85 for inputs exceeding 64 spikes (Fig. 2f). This relationship persisted after controlling for trial numbers and across levels of feature removal (Supplementary Fig. 2d). Thus, perturbation detectability increased systematically with the number of evoked target spikes.

Finally, the representation was transient. When ensemble identity was decoded independently from each 22-ms imaging frame, decoding was concentrated around frames overlapping the 100-ms stimulation period and peaked within this window for all ensemble sizes (Fig. 2g; Supplementary Fig. 4). Peak accuracy increased from ∼20% for single targets to ∼90% for 16-cell ensembles. The surrounding network therefore transiently preserves both the occurrence and identity of a local perturbation, with larger ensembles producing increasingly robust—but not disproportionately amplified—population representations.

### Responders and decoding errors are organized within ∼100 µm of the targets

We next examined the spatial organization of ensemble responses. The fraction of significant responders was highest near the stimulated targets (for multi-target ensembles, distance to all targets was included), reaching ∼0.6–0.8 within 50 µm at α = 1%, and declined with distance to a plateau by ∼150 µm that persisted to the edge of the field of view (Fig. 3a). Increasing ensemble size raised this plateau monotonically, from ∼0.15 for single targets and pairs to ∼0.3 for 16-cell ensembles, without substantially changing the spatial extent of the response. Thus, larger ensembles recruited a greater fraction of neurons throughout the network rather than expanding the radius of local recruitment. This spatial profile was preserved across significance thresholds from α = 1% to 40%; the NonR fraction showed the complementary pattern, whereas NegR became prominent only at more permissive thresholds (Supplementary Fig. 5). Stimulation of all 32 targets in a field of view extended the same trend (Fig. 3a; Supplementary Fig. 5).

The same spatial scale also structured errors in decoding ensemble identity. We calculated confusion matrices for the identity classifiers and asked whether targets that were more frequently confused were also physically closer. In an example dataset, pairwise confusion was ∼10% for targets separated by less than ∼30 µm but fell to ∼1% beyond 100 µm (Fig. 3b). Across all datasets, confusion similarly declined from ∼5% at the shortest separations to ∼2% beyond ∼100 µm (Fig. 3c). Pairs of 2-cell ensembles showed the same relationship when distance was measured between ensemble centres of mass (Fig. 3c, inset).

Thus, the spatial organization of decoding mirrored that of network recruitment. Nearby targets generated more overlapping population responses and were therefore more likely to be confused, whereas targets separated by more than ∼100 µm were represented by increasingly distinct response patterns. This distance closely matches the spatial scale over which local excitatory connectivity declines in superficial cortex [1–5,38].

### Ensemble-size normalized responses collapse onto a common baseline-dependence curve

In our previous single-neuron experiments, responses to evoked spikes were largely independent of preceding network activity at low baseline levels, but increased with baseline activity above a threshold [18]. The ensemble ladder allowed us to test whether the effects of input strength and ongoing network state could be separated.

For each ensemble size, the mean evoked spike count per non-target neuron was approximately independent of pre-stimulus baseline activity below a certain level and increased with baseline activity above it (Supplementary Fig. 6). When plotted together, the response curves were naturally vertically ordered by ensemble size (Fig. 4a). Normalizing each response by ensemble size raised to the power 0.285—the exponent governing the ensemble input–response relationship—collapsed the five curves onto a single function and minimized the collapse error (Fig. 4b, inset).

**Fig. 4.**
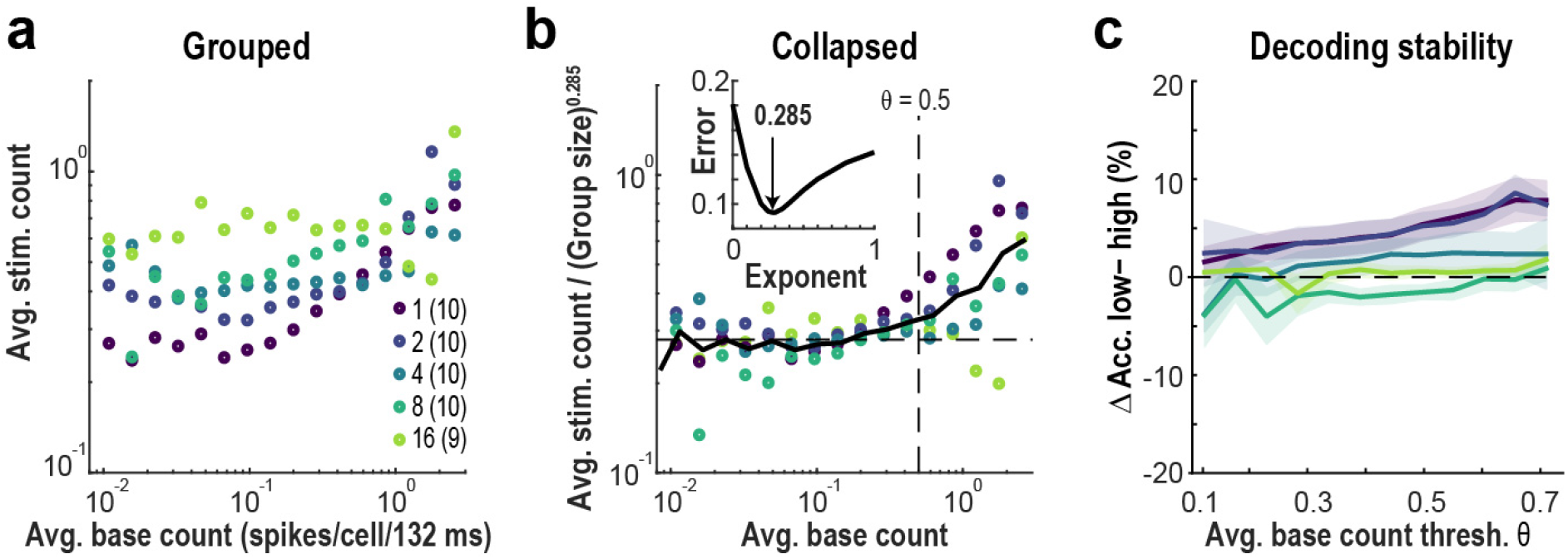
Ensemble-size normalization collapses responses onto a common baseline-dependence curve. **a,** Mean evoked spike count per non-target neuron versus preceding baseline activity for ensemble sizes 1, 2, 4, 8 and 16 (circles, binned means; log–log axes). **b,** Same data normalized by ensemble size to the power of 0.285. Horizontal dashed line, common response floor; diagonal dashed line, slope 0.5 for reference; solid line, average across normalized data. Inset, collapse error (standard deviation over the average across normalized data) as a function of normalization exponent, with a minimum at 0.285. Per-ensemble-size distributions and variability are shown in Supplementary Fig. 6. c, Difference in decoding accuracy between trials with Low (<θ) and High (≥θ) base activity for ensemble sizes 1–16. Shading, mean ± s.e.m..

The collapsed relationship revealed two regimes. Below a baseline activity of ∼0.18 spikes per 132 ms, responses remained at a floor of ∼0.27 spikes per 132 ms and were largely independent of ongoing activity. Above this level, responses increased with baseline activity (Fig. 4b, right). Decoding accuracy did not differ significantly between the low- and high-baseline activity regimes (Fig. 4c).

Thus, the effects of ensemble size and ongoing cortical activity were approximately separable: ensemble size scaled the magnitude of the response through the same ∼0.285 exponent that governed input–response scaling, whereas baseline network activity modulated the response through a common state-dependent function.

## Discussion

Using all-optical interrogation of L2/3 in awake mouse visual cortex, we asked how the network transforms the coordinated activity of one to sixteen pyramidal neurons. Across nearly two orders of magnitude in evoked spike count, ensemble responses followed the same power law with exponent ∼0.285 previously observed for single-neuron perturbations [18]. We found no threshold at which co-activation produced disproportionate amplification: responder recruitment was sublinear, suppression increased with ensemble size, and ongoing activity retained its parabolic avalanche structure. Yet the identity of the stimulated ensemble remained decodable from the surrounding population, with accuracy and robustness increasing with ensemble size. Thus, larger ensembles produce stronger and more distinguishable responses without entering a qualitatively different propagation regime.

This result challenges the idea that coincident activity must acquire special collective efficacy to overcome the sparse and weak connectivity of superficial cortex [1–9,12–14]. If synchrony generated cooperative amplification, ensemble responses should deviate upward from those predicted by their spike count. Instead, response magnitude varied smoothly with evoked input, and normalizing by ensemble size to the power of 0.285 collapsed responses onto a common baseline-dependence curve. Within the range tested, increasing the number of co-active neurons moved the population response along the same compressive input–output function observed for weaker perturbations. This does not diminish the behavioural importance of single neurons or ensembles. Single-neuron perturbations have nonetheless revealed measurable effects on neighbouring neurons and on behaviour [39–41]. Similarly, small groups of cortical neurons can influence perception or recall learned behaviour [15–17]; our results suggest that such effects need not require a circuit-level transition [42,43]. Increasing ensemble activity may simply move the network continuously along a compressive response curve until a downstream behavioural threshold is crossed.

The sublinear recruitment of responders likely reflects two related features of local cortical circuitry. Nearby targets recruit overlapping responder populations, as indicated by the spatial clustering of responders and by decoding errors between targets separated by less than ∼100 µm. At the same time, suppression increased with ensemble size, consistent with coordinated recruitment of excitation and inhibition in balanced cortical networks [33,34,44–46]. Together, overlapping excitation and increasing inhibition can limit the growth of the recruited population while preserving the shape of the network response function. The shortfall from additive responder recruitment is therefore the cellular counterpart of the sublinear scaling observed in spike counts.

The recurrence of an exponent near 0.285 across single-neuron and ensemble perturbations supports the interpretation that local cortical propagation is governed by near-critical dynamics. This value is expected for a two-dimensional system near a directed-percolation transition [25,26], consistent with the sheet-like organization of superficial cortex for local L2/3 interactions [47–49]. Such compressive scaling expands dynamic range [28], allowing the same network to represent inputs spanning orders of magnitude without saturation in line with the psychophysical compression described by Stevens’ law [26,50]. The ensemble ladder extends this relationship far beyond the input range available with single-neuron stimulation. Moreover, the collapse of responses across baseline activity suggests that input strength and network state make approximately separable contributions: ensemble size determines how far the response moves along the critical scaling function, whereas ongoing activity modulates the gain with which that input is expressed [18,51,52].

The network nevertheless retained detailed information about which neurons had fired. Ensemble identity was decodable at every size, became more robust as ensemble input increased, and remained distributed across the population despite sparse classifier evidence on individual trials. Detectability increased systematically with the number of evoked spikes, consistent with larger ensembles improving coding by recruiting more of the network rather than by engaging a new coding mechanism. Decoding confusion was concentrated over ∼50–100 µm, matching the spatial scale of local excitatory connectivity and responder clustering [1–5,38], while more distant targets remained distinguishable despite overlapping population representations [53,54]. Together, these findings separate two consequences of coordinated firing: coordinated firing strengthens identity without gaining disproportionate causal influence. Near criticality, cortical networks preserve input identity while transforming input magnitude through a common scale-invariant response function.

## Methods

### Experimental design and analysis units

The dataset comprised 10 imaging recording sets from 9 adult mice (animal sex was not considered in the study analysis due to the low number of animals included.). Each recording sampled one V1 field of view containing a hierarchy of 32 target cells tested principally as ensembles of 1, 2, 4, 8 and 16 neurons. A 32-cell condition was acquired in 2 of the 10 recordings and is reported only where indicated. The same field of view and target hierarchy were followed across the sequential stimulation sessions. When data was pooled, statistics was reported average across the 10 recording sets.

### Animal surgery

All procedures were approved by the NIH Animal Care and Use Committee (ACUC) and experiments followed the NIH Guide for the Care and Use of Laboratory Animals. Transgenic Emx1-Cre mice were obtained from Jackson labs (RRID:IMSR_JAX:005628), bred inhouse with C57BL/6 backgrounds (Jackson Laboratory) under a reversed 12:12 h light/dark cycle. Chronic 2PI in adult (>6 weeks) mice was enabled by using a head bar in combination with a cranial window placed centered at ∼2.5 mm from the midline (right hemisphere) and ∼1 mm rostral to the lambdoid suture and consisting of a stack of circular glass cover slips using established protocols [55]. Mice were injected with a viral construct to express jGCaMP8s [56] and ChrimsonR [57] in pyramidal neurons [58]. A total of 2–3 injections of virus (100–400 nL; <1 µL in total; 10^13 vg/mL; AAV9-hSyn-DIO-jGCaMP8s-P2A-stChrimsonR, Addgene) were administered into the right hemisphere. Chronic 2PI started after >2 weeks in identified V1 at a depth of ∼100–200 µm. To prepare for retinotopic mapping and 2PI imaging and stimulation, mice were briefly anesthetized (isoflurane ∼2%, <2 min) and then quickly moved over to the recording platform. After fixing the animal’s head using a head bar mount, the mouse was allowed to recover from anesthesia for ∼30 min before recordings started.

### Identification of V1 maps

Retinotopic maps of V1 and higher visual areas (HVAs) were generated for all mice prior to recording using published protocols [59,60]. Briefly, awake, head-fixed mice faced with their left eye a 19" LCD monitor placed at 10 cm distance and tilted 30° towards the mouse’s midline. Using Psychophysics toolbox [61], contrast-reversing, spherically corrected checkerboard bars were drifted across the screen vertically (altitude) and horizontally (azimuth) for each of the four directions (30 repeats per direction). Simultaneous wide-field imaging (Quantalux, Thorlabs) captured calcium indicator fluorescence, which was averaged for each direction. Altitude and azimuth phase maps were calculated by phase-wrapping the first harmonics of the 1D Fourier transform for each of the four averages and subsequently subtracting the maps of the opposite directions [60]. Sign maps were generated by taking the sine of the angle between the gradients in the altitude and azimuth maps and processed [59]. Borders were drawn around visual area patches and overlaid onto anatomical reference images to identify V1.

### Visual stimulation and orientation preference

Before and after the holographic stimulation series, large-field drifting gratings of 8 orientations were presented to the left eye (400 trials, 1 s stimulus duration, 1 s stimulus interval, 100% contrast, 30 degrees per cycle at 4 Hz; Fig. 1b). For each imaged neuron, visual responsiveness and preferred orientation were determined from the trial-averaged spike counts during grating presentation by fitting a double Gaussian to its visual-stimulus response curve. Visual responsiveness was defined based on goodness-of-fit (> 0.7) and orientation selectivity index (> 0.3). The difference in preferred orientation between a non-target neuron and its target was computed as the smallest angle between their preferred orientations (0–90°). All analyses were based on the single-target ensemble recordings.

### Holographic stimulation of neuronal ensembles

Holographic stimulation [62] was done by manually selecting individual pyramidal neurons as targets from the 2-photon image of the field of view. All neurons that expressed jGCaMP8s were presumed to also express ChrimsonR (due to the P2A virus strategy) and were therefore available for being selected for stimulation. Thirty-two targets were selected per field of view, avoiding cells close to the edges. A map of top-hat patterns with diameter of 10 µm was used to generate the hologram on the Spatial Light Modulator (SLM, Meadowlark, LCoS high resolution 1920×1152 phase modulator). Light from the laser/OPA system (Carbide/Orpheus-F, Light Conversion; λ = 1064 nm, repetition rate = 600 kHz, pulse width <100 fs) was expanded (6x) to fill the area of the SLM (8 mm × 8 mm). The hologram on the pupil plane was shrunk to fit the size of the Galvo-Galvo scanners (∼3 mm diameter) and form the desired pattern of top hats after the objective lens (Nikon 16X, working distance = 3 mm and effective focal length = 12.5 mm) on the selected targets. For ensemble stimulation, a single hologram addressed all targets within the ensemble simultaneously, with ∼5–10 mW per target in the ensemble (∼5–160 mW typical total power; up to ∼320 mW on 2 experiments in which all 32 targets were stimulated simultaneously). No artifact from this stimulation was detectable in our imaging with our combination of indicator and opsin, stimulation wavelength and the low power required, as documented previously [18].

Ensembles were formed hierarchically. The 32 single targets were merged into 16 pairs by joining spatially neighbouring targets; pairs were merged into 8 quartets, quartets into 4 octets, and octets into 2 ensembles of 16, again by spatial proximity, so that each group at one level was the union of two groups at the previous level (Fig. 1b). Each ensemble size was recorded in a separate session of ∼17 min in the same field of view, in the fixed order 1, 2, 4, 8, 16. In 2 out of the 10 recordings all 32 targets were stimulated as one ensemble after the 16-targets ensemble recording. Within a session, the groups of a given size were stimulated in a pseudo-random sequence, each group for 100 ms per trial and 100 trials per group, with a fixed inter-stimulus interval (onset-to-onset) per ensemble size of 0.3, 0.6, 1.2, 2.4 and 4.8 s for ensembles of 1, 2, 4, 8 and 16 neurons, respectively, so that the total stimulation burden/session duration remains the same for each recording. No photobleaching, or other artifacts were observed as a consequence of the chosen ensemble-size order for the recordings.

### 2PI, pre-processing pipeline and meta data collection

For standard 2PI, images were acquired by a scanning microscope (Bergamo II series, B248, Thorlabs Inc.) coupled to a pulsed femtosecond Ti:Sapphire 2-photon laser with dispersion compensation (Chameleon Discovery NX, Coherent Inc.). The microscope was controlled by ThorImageLS and ThorSync software (Thorlabs Inc.). The wavelength was tuned to 940 nm to excite the calcium indicator jGCaMP8s. Signals were collected through a 16× 0.8 NA microscope objective (Nikon). Emitted photons were collected through a 525/50 nm band filter using GaAsP photomultiplier tubes. The field of view was ∼450×450 µm². Imaging frames of 512×512 pixels were acquired at 45.527 Hz by bidirectional scanning of a 12 kHz Galvo-resonant scanner. Beam turnarounds at the edges of the image were blanked with a Pockels cell. The average power for imaging was <70 mW, measured at the sample.

The obtained tif-movies in uint16 format were rigid motion-corrected via the python-based software package ‘suite2p’ [63]. Registered images were further denoised using machine-learning based, deep interpolation [64] (see below) and then semi-automatically processed by suite2p for regions of interest (ROI) selection and fluorescence signal extraction. We performed visual curation of the automatically selected ROIs to ensure only neurons were included in the analysis. Neuron identity was tracked across the sequential recordings of one field of view using the ROIMatch algorithm (https://github.com/ransona/ROIMatchPub; average of ∼98 neurons matched per experiment across 7 recordings). For each labeled neuron, raw soma and neuropil fluorescence signals over time were extracted. Spiking probabilities were obtained from neuropil-corrected fluorescence traces (Fcorrected = FROI – 0.7*Fneuropil) via MLspike (https://github.com/MLspike), utilizing its autocalibration feature to obtain unitary spike event amplitude, decay time, and channel noise for individual neurons.

#### Deep-interpolation

Deep-interpolation [64] (Deep-IP; https://github.com/AllenInstitute/deepinterpolation) removes independent noise by using local spatiotemporal data across a noisy image stack of Npre + Npost frames to predict, or interpolate, pixel intensity values throughout a single withheld central frame. The deep neural network is a nonlinear interpolation model based on a UNet inspired encoder-decoder architecture with 2D convolutional layers where training and validation are performed on noisy images without the need for ground truth data. As described previously in detail [18,19], after rigid motion correction, individual denoised frames were obtained by streaming one 60-frame (Npre = Npost = 30 frames) registered, image stack through the provided Ai-93 pretrained model [64] for each frame to be interpolated. At an imaging rate of ∼45 Hz, these 60 frames correspond to a combined ∼1.3 seconds of data surrounding the frame to be interpolated.

### Postprocessing pipeline

Our postprocessing pipelines were custom-written in Matlab (Mathworks) and Python (www.python.org). Some routines utilized NumPy (https://numpy.org/) and Matplotlib (https://matplotlib.org/).

#### Stimulus response measurement

Unless stated otherwise, spike count over a window of 6 imaging frames (∼132 ms) starting at the stimulation onset was used as a measure of the response to holographic stimulation for all cells. That window includes the full stimulus duration (100 ms) and one extra frame after the stimulus offset, for which increased activity in the targets can still be observed (Fig. 1d). This was compared to the baseline count, defined as a 6-frame window starting ∼182 ms before stimulus onset. The evoked ensemble spike count of a trial was defined as the sum of the response-window spike counts over all members of the stimulated ensemble.

#### Ensemble selection and exclusion

Ensembles that failed to respond to holographical stimulation were excluded from analysis using the following criteria. For each target, a minimum evoke spike count threshold was defined as a count above the 90th percentile of the baseline count. Then, each trial we stimulated a given ensemble was classified as a fully responsive trial (when all targets in the ensemble had evoked spike count above the threshold), partially responsive trial (when at least one of the targets in the ensemble had evoked spike count above the threshold) or not responsive trial (when none of the targets in the ensemble had evoke spike count above the threshold). Ensembles that had a fraction of not responsive trials above 20% were excluded from the analysis. The number of excluded ensembles are: 112/320 (35%; 1-target ensembles), 24/160 (15%; 2-target ensembles), 1/80 (1.25%; 4-target ensembles) and 0 for ensembles of size 8+. For decoding analyses, trial and target filtering are described below.

#### Definition of significant responders

Cells with a significant response to the stimulation of a group were defined as those whose response spike count distribution differed significantly from their baseline count distribution (z-test, α = 2.5% unless stated otherwise; Fig. 3a and Supplementary Fig. 5 use α = 1% to 40%). A cell was defined as a positive responder (PosR) if its response count was significantly larger than baseline, a negative responder (NegR) if significantly smaller, and a non-significant responder (NonR) otherwise. This procedure was applied separately to every group of every ensemble size, so that a non-target cell could be a significant responder to some groups and not to others. Given the low baseline activity of most cells, negative responders are likely underestimated with spike data [18].

#### Expected number of responders

For an ensemble at a given level of the hierarchy, the expected number of PosR under the assumption that each target contributes its own responder pool was computed as the size of the union of the PosR sets obtained for the constituent single targets in the single-target recording, at the same significance threshold. The difference between the measured number of PosR of the ensemble and this expectation was averaged over groups and datasets (Fig. 2b).

#### Correlation analysis

Pairwise correlations of the complete spike trains were computed between target neurons of the same 2-target ensemble, between targets and the non-target population, and between control pairs of all non-target neurons, for the reference recordings before and after the series and for each ensemble size, and normalized to (divided by) the control value. Values are reported as mean ± SE across experiments. Significance of correlation changes across conditions was obtained by employing the Wilcoxon rank sum test at a significance level of 0.05.

#### Response scaling and ensemble-size collapse

Response curves (Fig. 2c, d; Supplementary Fig. 2a, b) were obtained by binning trials by the evoked ensemble spike count (or by ensemble size, or by the number of active targets) and averaging the response-window spike count of PosR, NonR or all non-target neurons within each bin. Power-law exponents were obtained by linear regression in log–log coordinates; best fits and 95% confidence intervals are reported in Supplementary Table 1. For the ensemble-size collapse (Fig. 4b), trials of each ensemble size were binned by the mean baseline spike count preceding the trial, and the mean evoked spike count in each bin was divided by the ensemble size to the power of 0.285 (Fig. 4b); the per-size curves are shown in Supplementary Fig. 6. Collapse error was calculated as the average standard deviation of the mean curve (average over the 5 different ensemble sizes) as function of the collapse exponent.

### Decoding analyses

#### Decoding of stimulated-ensemble identity

As in our previous work [18], the decoding analyses were performed separately for each dataset (n = 10), the main difference in this study being that for each dataset we also examined stimulation of multiple neurons at the same time, each targeted ensemble now being treated as a decodable class. We examined ensembles with 1, 2, 4, 8 or 16 targeted neurons, corresponding to 32, 16, 8, 4 and 2 distinct classes in our decoding paradigm. Spiking activity was binned at the imaging frame rate (≈45.5 Hz, 22-ms bins). We treated every stimulation trial as an independent sample and the individual neuron responses as features. The values of these features were obtained as the mean spike count of a given feature neuron over a predefined time window, expressed in terms of the number of frames, nw. We typically used nw = 6 frames following stimulation onset (0–132 ms), a window that covers the 100 ms optogenetic stimulation window and its immediate aftermath. In other cases, windows of size nw = 1, 2, 4 and 8 were used to explore their effect on accuracy; in particular, we used single-frame activity as a feature when running time-resolved decoding to increase the time resolution (see below). The feature vector contained only non-targeted neurons: all neurons belonging to any stimulation ensemble were excluded from the features, and trials in which the targeted cells could not be identified in the imaging data were discarded. The classification label was the index of the stimulated ensemble, so chance performance equals 100%/nc for nc classes.

Unless stated otherwise, the classifier was a gradient-boosted decision-tree ensemble (XGBoost [35]) with 1,000 trees and otherwise default hyperparameters. Performance was estimated over 100 random stratified train/test splits (80%/20%); we report the mean percent accuracy on the held-out test set, with shaded bands giving the standard error of the mean (SEM) across splits or, for dataset averages, across datasets. All random splits and model seeds were derived deterministically from a base seed, making the analysis reproducible from a fixed base seed. Where indicated, an optional trial filter retained only trials in which the stimulated ensemble responded (at least one validly identified ensemble neuron active in the response window) and dropped classes with fewer than 20 responsive trials; the analyses shown in Figs. 2 and 3 use the filtered data, and the unfiltered data are shown in Supplementary Fig. 2c.

#### Neuron importance ranking and feature-removal curves

To quantify how decodable information is distributed over the non-targeted population, neurons were ranked by their importance for the trained classifier using Shapley additive explanations (SHAP, TreeExplainer [37]). For each of 50 stratified train/test splits, a classifier was trained and the importance of each neuron was computed as the mean absolute SHAP value over all samples and classes; importances were then averaged over splits to give a single best-coding ranking per dataset and ensemble-size condition. A complementary leave-one-out ranking (the change in decoding accuracy when a single neuron was omitted, averaged over 100 splits) produced qualitatively similar removal curves (Supplementary Fig. 2c).

Feature-removal curves were obtained by removing the top-ranked (best-coding) neurons cumulatively and retraining the classifier from scratch on the remaining neurons at each removal step, again over 100 stratified train/test splits per step. Because datasets differ in neuron counts, each dataset’s curve was expressed as accuracy versus the percentage of neurons removed, linearly interpolated onto a common 0–100% axis, and averaged across datasets (Fig. 2e; n = 10 datasets per condition, n = 9 for the 16-cell condition, for which one dataset did not yield a valid decoding problem). Shaded bands show the SEM across datasets; dashed lines mark chance (1/nc).

#### Detection of stimulation from pooled evoked spike counts

To ask whether the mere occurrence of stimulation can be read out from the non-targeted population and how this depends on the evoked drive, we performed a binary classification of stimulation versus baseline for which we could pool the data from different ensembles. The reason we chose binary classification problem is that when stratifying all trials based on the number of activated target neurons, performing multi-class decoding would again subdivide our data into smaller chunks. The positive examples were the response-window activity vectors (non-targeted neurons) of stimulation trials and they were matched with an equal number of negative examples, drawn from baseline activity in windows shifted to precede stimulation onset in the same recordings. Trials were grouped by the total number of spikes fired by the targeted ensemble in the response window, pooled into bins of 1, 2, 3–4, 5–8, 9–16, 17–32, 33–64 and >64 evoked spikes, and a separate classifier (XGBoost, default settings) was trained for each pool.

Performance was measured as the area under the ROC curve (AUC) on a held-out test set. Feature removal here used recursive SHAP-guided elimination: at every step the model was retrained, the neuron with the largest mean absolute SHAP value was removed, and the test AUC was recorded, until no features remained. Curves in Fig. 2f show the mean AUC (± SEM across datasets) as a function of the percentage of best-coding neurons removed; the inset shows the same data on a logarithmic axis for the higher spike-count pools. Detectability increased systematically with the total evoked spike count, and an equal-trial-number control confirmed that this ordering is not a sample-size artifact. The dependence of the AUC on evoked spike count at each level of feature removal is shown in Supplementary Fig. 2d.

#### Time-resolved decoding

To resolve when the ensemble identity is decodable, the same decoding procedure was applied to a sliding window moved across the recorded spike activity raster. For Fig. 2g the window was a single 22-ms frame advanced in steps of one frame, from 500 ms before stimulation onset to 1 s after; at each offset the classifier (XGBoost, 1,000 trees) was trained and tested on that window alone, using 50 stratified 80/20 train/test splits. Windows never mixed pre- and post-stimulus frames of neighbouring trials. Curves show, for each ensemble-size condition, the accuracy averaged over datasets as a function of window time relative to stimulation onset; horizontal dashed lines mark the corresponding chance levels, and the shaded vertical bar marks the 100-ms stimulation period. For the six-frame analysis (Supplementary Fig. 4, right column), the fading pink shading indicates that fewer and fewer frames from the stimulation window contribute to the averaged feature. Note that there is additional temporal smoothing due to our use of DeepInterpolation, an issue explored previously [18].

#### Distance dependence of decoding confusion

To test whether decoding errors reflect spatial proximity of the stimulated targets, full confusion matrices were estimated for all ensemble conditions, using the same features and classifier as above. Only for the 32-class case do distances reflect true pairwise Euclidean distances between targets; for ensembles, distances were estimated between ensemble centres of mass. Test-set confusion matrices were accumulated over 100 stratified train/test splits and row-normalized, so that entry Cij gives the percentage of trials of true class i classified as class j (Fig. 3b, inset; colour scale saturated above 30%). Off-diagonal confusion between a pair of classes, Cij and Cji, was symmetrized using their harmonic mean, 2·Cij·Cji/(Cij + Cji), and plotted against distance for an example dataset (Fig. 3b). Arithmetic and geometric means gave no significant differences, and harmonic-mean results are shown. The solid black line represents the mean confusion within consecutive fixed-width distance bins (bin size 15 pixels = 13.2 µm). Fig. 3c shows the same data pooling all pairs over all datasets, and the inset shows the same trend for 2-cell ensembles. For larger ensembles the centre-of-mass distance does not faithfully reflect the overlap between individual neurons and those data are not shown. The association between confusion and distance was quantified with the Spearman rank correlation over all pooled class pairs; confusion was elevated for targets closer than ≈50–100 µm and essentially flat beyond.

#### Single-neuron selectivity and per-trial importance

For each target, a selectivity index *SI* was computed for every non-target neuron as follows. Let *x_ij_* be the response of non-target neuron *j* on trial *i* (its mean spike count in the 132-ms window starting on stimulation), and *E* the set of trials in which ensemble *e* was stimulated:

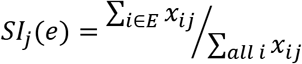

The range of *SI* is mathematically bound between 0 and 1, but for a completely unselective neuron with balanced ensembles, *SI* ≈ 1⁄*N_e_*, where *N_e_* is the number of ensembles, so the “chance” *SI* is always greater than 0. When *SI*(*e*) = 1, it means the neuron only fires when ensemble *e* is stimulated.

*SI* was then compared with the neuron’s mean absolute Shapley value for that target/ensemble (Supplementary Fig. 3a,b). Per-trial contributions were summarized by the entropy of the distribution of absolute Shapley values across neurons in each trial and displayed as neuron-by-trial maps (Supplementary Fig. 3c,d).

#### Software

All decoding analyses were implemented in Python 3 using NumPy, SciPy, scikit-learn [65], XGBoost [35] and the SHAP library [37], and were run on the NIH Biowulf high-performance computing cluster. Every analysis stage caches intermediate results (importance rankings, per-step accuracies, confusion matrices), making runs resumable and reproducible from a fixed base seed.

### Avalanche analysis

#### Continuous epochs of suprathreshold population activity

Continuous periods of population activity were identified by applying a threshold Θ on the population activity, the sum of the spikes from all non-target neurons at a given time, as described previously [18,19]. For a given recording and coarse-graining value k, the dependence of the number of epochs on the threshold, N(Θ), was obtained for a range of thresholds from one that removed no population activity to one that removed all population activity. N(Θ) was typically well approximated by a log-normal distribution, and the threshold used in the analysis for all recordings and coarse-graining factors was chosen as its mean, estimated for each k. See ref. 29 for details.

#### Temporal coarse-graining

A temporal coarse-graining operation was applied to the thresholded population activity. For a given temporal coarse-graining factor k, an ensemble of k coarse-grained time series was obtained by summing k consecutive frames at each of the k possible phase offsets [19]. For each time series, epochs were extracted as maximal contiguous runs of suprathreshold activity; the size of an epoch is the total number of spikes within it and its duration the number of coarse-grained bins multiplied by k times the frame duration. For a given ensemble of coarse-grained time series, all epochs were pooled. Avalanche size and duration distributions were computed separately for the reference recording before the series, for each ensemble size, and for the reference recording after the series (Supplementary Fig. 1).

#### Scaling curve fit

For more precise evaluation of the scaling curve, mean avalanche size versus duration, we used a double power law with initial slope χ_short, transitioning to a second slope χ_long at around a duration Φ, with a transition sharpness parameter fixed at 4 for all curves, as described previously [18]. All fits were performed in log-space.

### Statistics & Reproducibility

No statistical method was used to predetermine sample size. Experiments were not randomized, and the investigators were not blinded to allocation during experiments and outcome assessment. Animal sex was not considered in the study analysis due to the low number of animals included. For statistical analysis, please refer to each relevant Methods subsection for details on tests used and sample sizes. Schematic drawings and layouts were created using Adobe Illustrator v28.

## Data Availability

The pre-processed imaging data used in this study will be made available in Zenodo. Source data for all main and supplementary figures will be provided with the paper.

## Code Availability

Computer code used in this study will be made available on GitHub.

## Acknowledgements

We thank members of the Plenz lab for help with experiments, data analysis, animal surgery and care. We thank the members from our U19 BRAIN initiative grant for the many discussions and technical advice. This research was supported by the Division of the Intramural Research Program (DIRP) of the National Institute of Mental Health (NIMH), USA, ZIAMH002797, ZIAMH002971, and the BRAIN initiative Grant U19 NS107464-01. This research utilized the supercomputing resources of the National Institutes of Health (NIH, USA; Biowulf, http://hpc.nih.gov). Claude Code Opus 4.8 – 5.5 was utilized to assemble parts of the manuscript.

This research was supported in part by the Intramural Research Program of the National Institutes of Health (NIH). The contributions of the NIH author(s) are considered Works of the United States Government. The findings and conclusions presented in this paper are those of the author(s) and do not necessarily reflect the views of the NIH or the U.S. Department of Health and Human Services.

## Author Contributions

TLR and DP conceived and planned the study; TLR and BG performed experiments. TLR took the lead in data analysis. SP and RS took the lead in machine learning-based decoding of experimental data. All authors contributed to the analyses. TLR, SP and DP wrote the manuscript.

## Supplementary Figure Legends

**Supplementary Fig. 1.**
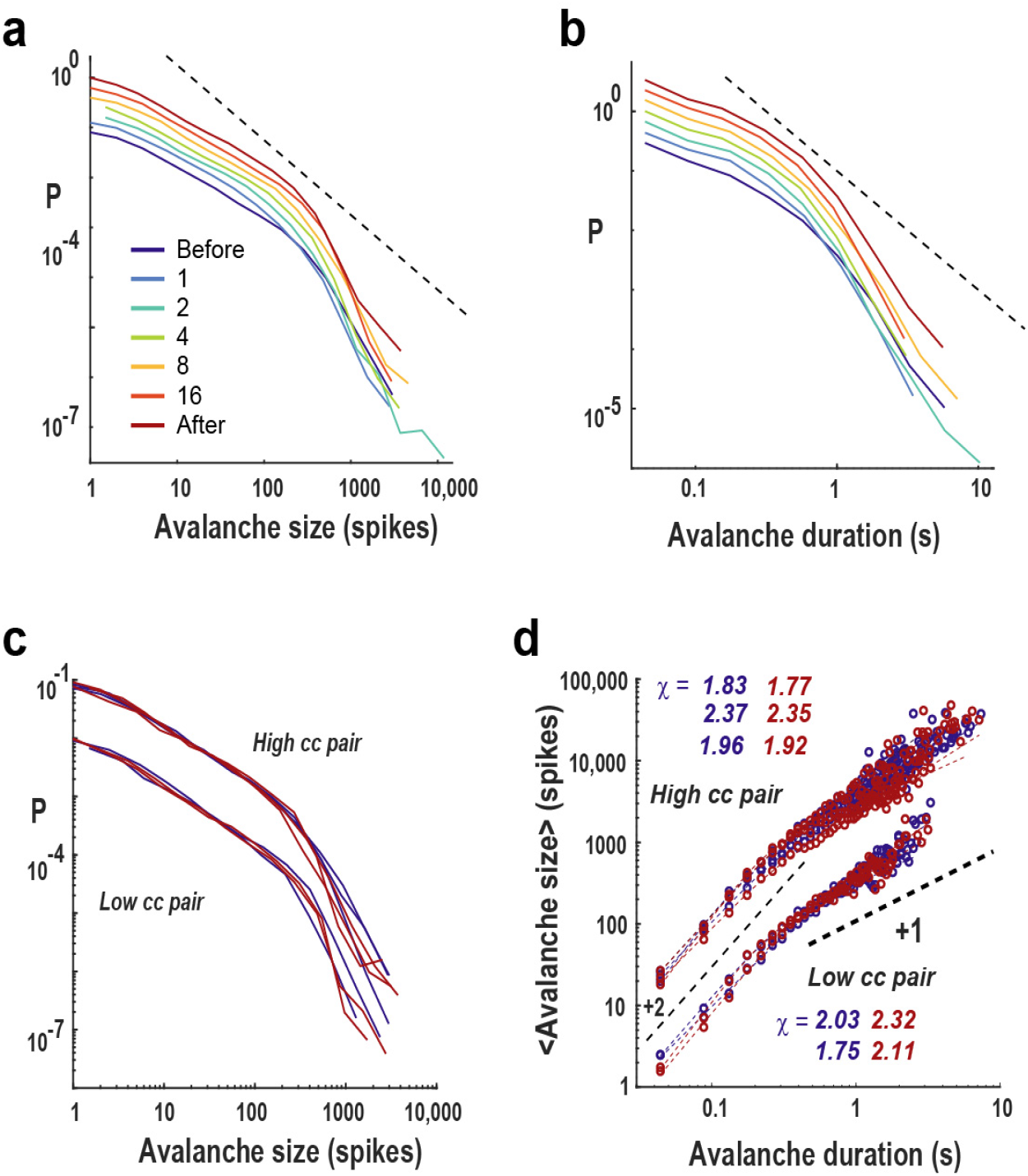
Avalanche statistics remain stable across ensemble stimulation. **a,b,** Avalanche size (a) and duration (b) distributions before, during stimulation of ensembles of 1–16 neurons, and after the stimulation series; curves are vertically offset for clarity. Dashed lines indicate reference power laws. **c,d,** Avalanche size distributions (c) and mean size–duration scaling (d) for high- and low-correlation target pairs before and after stimulation. Dashed lines indicate slopes of 2 and 1; fitted scaling exponents are indicated in the panels.

**Supplementary Fig. 2.**
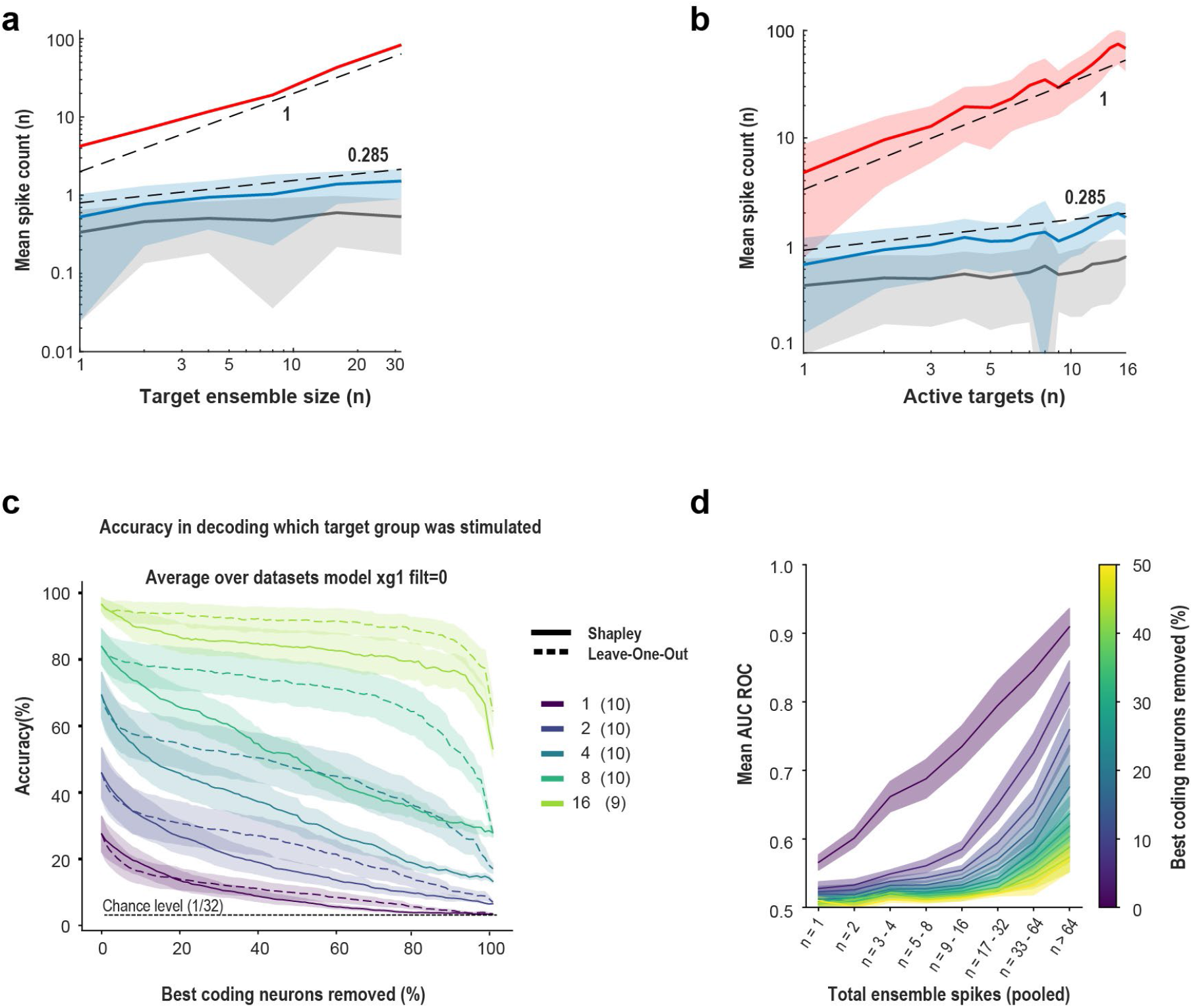
Response-scaling and decoding controls. **a,** Mean response-window spike count versus ensemble size for targets, PosR and NonR. Dashed lines indicate slopes of 1 and 0.285; shading, s.d. **b,** Same analysis versus the number of active targets. **c,** Ensemble-identity decoding during cumulative feature removal using Shapley-value or leave-one-out rankings without trial filtering. Shading, s.e.m. across datasets. **d,** Stimulation-detection AUC versus evoked target spike count at increasing levels of feature removal. Shading, s.e.m. across datasets.

**Supplementary Fig. 3.**
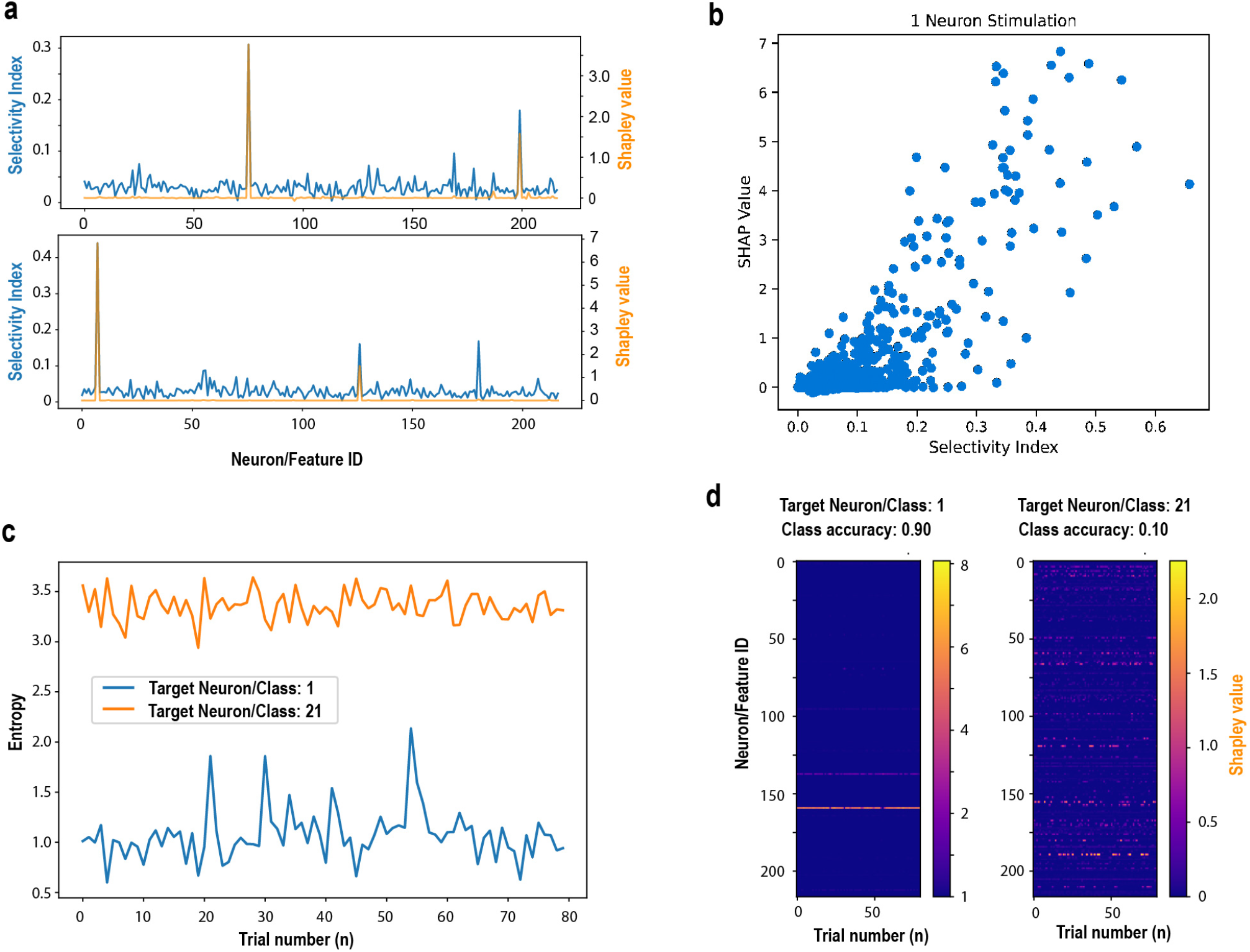
Classifier importance and neuronal selectivity. **a,** Selectivity index and mean absolute Shapley value for non-target neurons for two example single-cell targets. **b,** Shapley value versus selectivity across neuron–target pairs. **c,** Normalized entropy of per-trial Shapley contributions for well- and poorly decoded targets. **d,** Corresponding neuron-by-trial Shapley contributions. Selectivity and entropy are defined in Methods.

**Supplementary Fig. 4.**
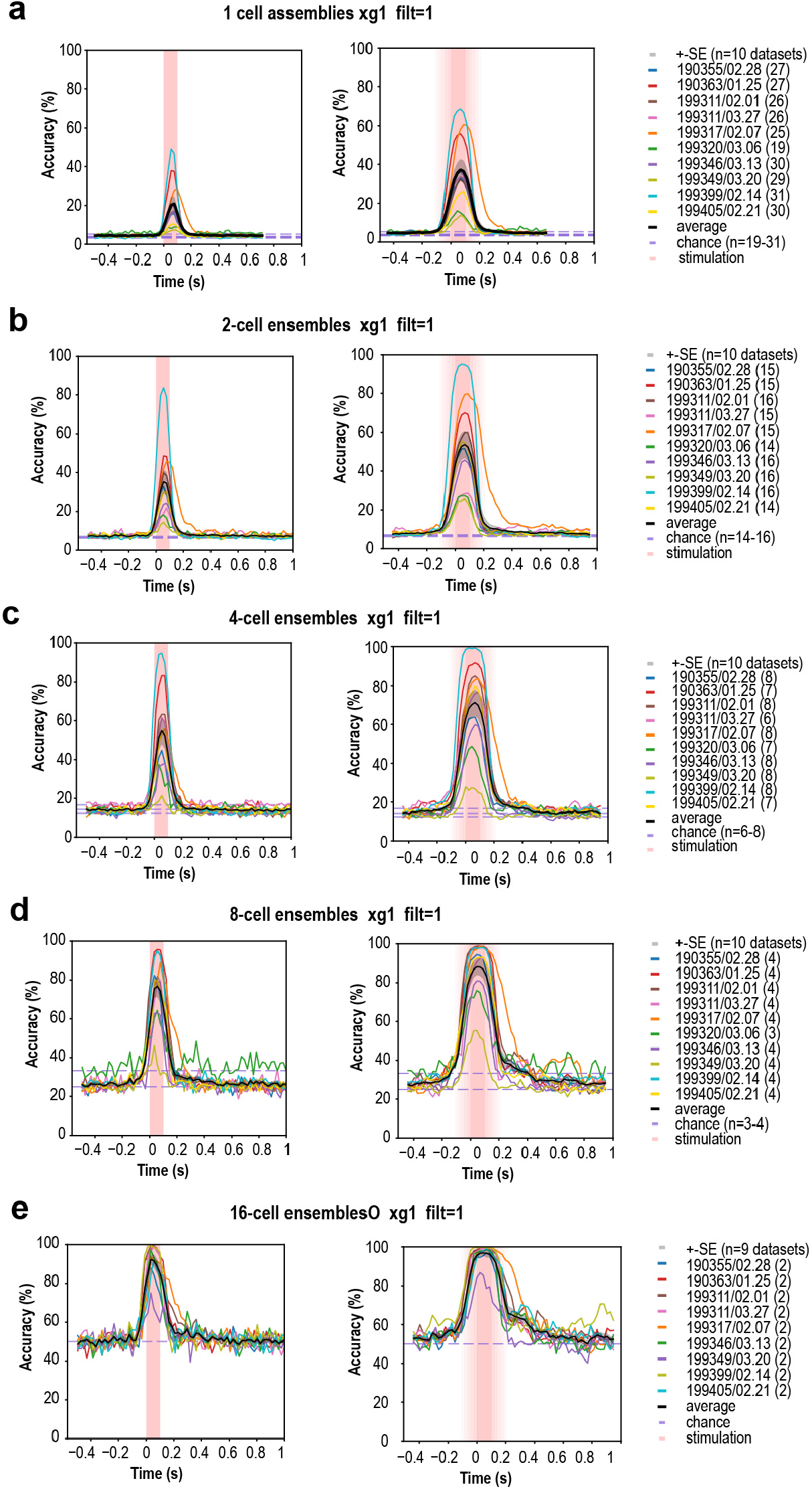
Time-resolved decoding across datasets. **a–e,** Ensemble-identity decoding versus time for ensembles of 1, 2, 4, 8 and 16 cells. Left, 22-ms decoding windows; right, 132-ms windows. Thin lines, individual datasets; thick line, mean; shading, s.e.m.; dashed lines, chance; pink shading, stimulation contribution to the decoding window. n = 10 datasets, except n = 9 for 16-cell ensembles.

**Supplementary Fig. 5.**
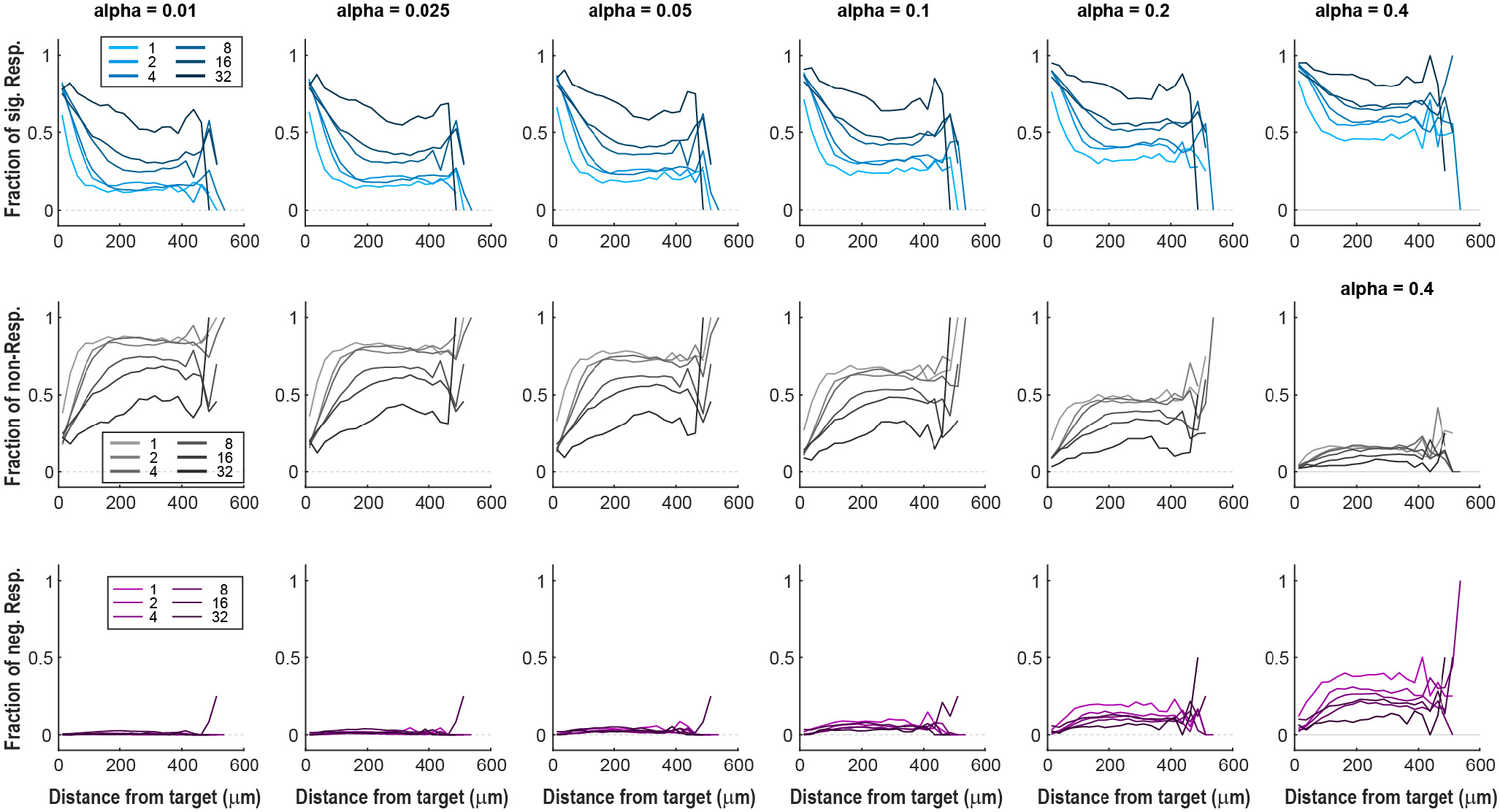
Responder-distance profiles across significance thresholds. Fractions of PosR, NonR and NegR versus distance from stimulated targets for ensemble sizes 1–32 (for multi-target ensembles, distance to all targets included in the analysis) and responder thresholds α = 0.01–0.4. The 32-cell condition was acquired in 2 of 10 recordings.

**Supplementary Fig. 6.**
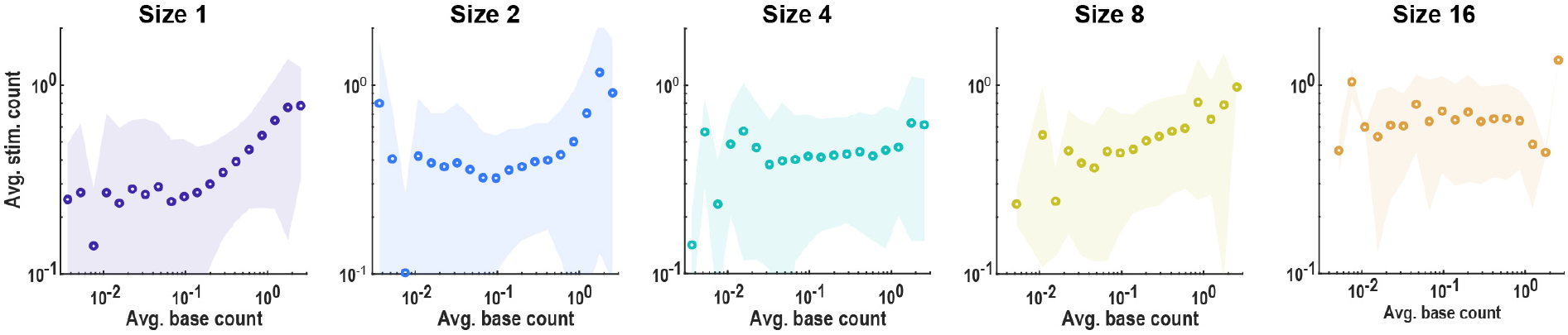
Baseline dependence for individual ensemble sizes. Mean response-window spike count per non-target neuron versus preceding baseline activity for ensembles of 1, 2, 4, 8 and 16 cells. Circles, binned means; shading, s.d.; vertical dashed line, baseline threshold of 0.18 spikes per response window. These data underlie the collapse in Fig. 4.

**Supplementary Table 1.**
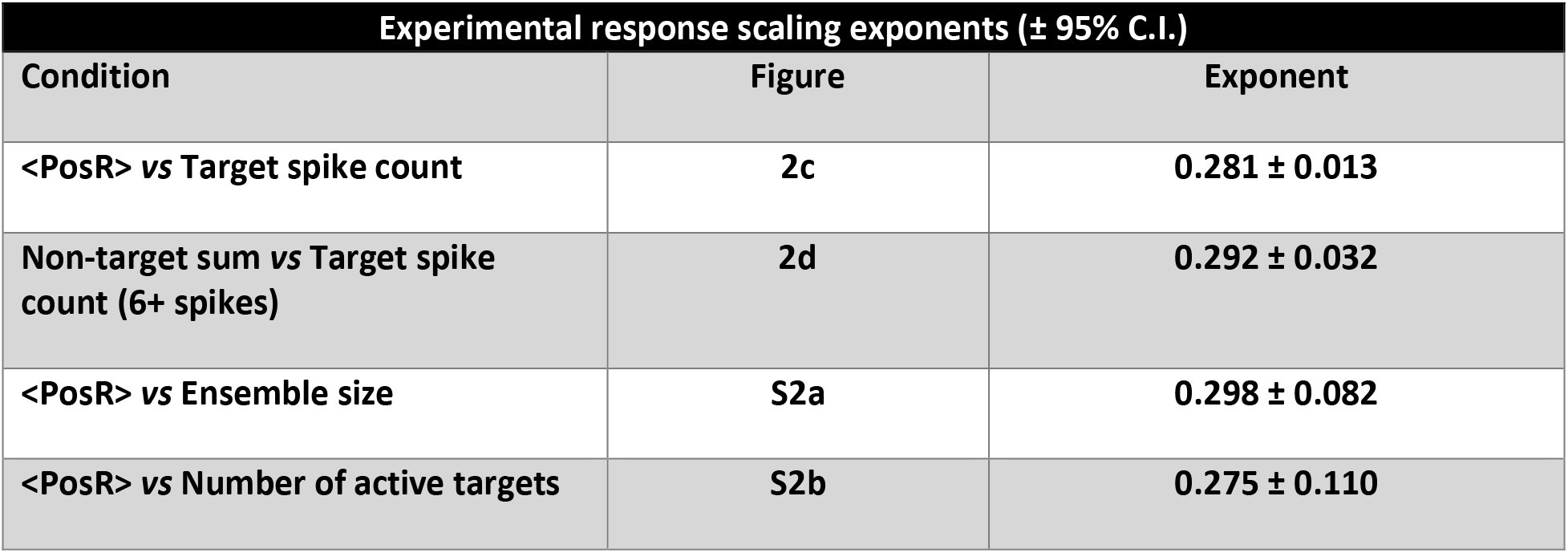
Response scaling exponents. Best-fit exponents and 95% confidence intervals for PosR, NonR and all-cell responses versus evoked ensemble spike count, ensemble size and the number of active targets, reported per dataset and for the pooled or hierarchical estimate.

| Experimental response scaling exponents (± 95% C.I.) | | |
| --- | --- | --- |
| Condition | Figure | Exponent |
| <PosR> vs Target spike count | 2c | $0.281 \pm 0.013$ |
| Non-target sum vs Target spike count (6+ spikes) | 2d | $0.292 \pm 0.032$ |
| <PosR> vs Ensemble size | S2a | $0.298 \pm 0.082$ |
| <PosR> vs Number of active targets | S2b | $0.275 \pm 0.110$ |

